# The turn less taken: Investigating patterns in *β*-turn dynamics using large-scale molecular dynamics data

**DOI:** 10.64898/2026.05.07.721674

**Authors:** Shuqin Zhang, Advaith Maddipatla, Sanketh Vedula, Alex M. Bronstein, Ailie Marx

## Abstract

*β*-turns are among the most common structural motifs in proteins, yet their conformational dynamics and sequence determinants remain incompletely understood. Here we present a data-driven classification and dynamic analysis of *β*-turn conformations using large-scale molecular dynamics trajectories from the mdCATH database. Clustering of backbone dihedral angles using a cross-bond Ramachandran representation identifies six *β*-turn types, including a previously uncharacterized hybrid I/I′ cluster that combines geometric features of canonical type I and I′ conformations. Time-resolved analysis indicates that this hybrid state acts as a transient intermediate state of *β*-turns. Transitions observed in molecular dynamics simulations closely match NMR ensembles and altlocs detected in X-ray crystal structures, with the most dominant exchanges occurring between type I and II, and between type I′ and II′ turns. Sequence analysis shows that each turn type exhibits characteristic amino acid preferences at the central residues (*i* + 1 and *i* + 2). Within these overall preferences, specific residue pairs display distinct biases toward static or dynamic behavior. Targeted in silico substitutions that interchange dynamic- and static-enriched residue pairs shift the conformational behavior of turns accordingly, providing direct support for these sequence-dynamics relationships. Analysis of flanking secondary-structure environments reveals that structural context further modulates turn flexibility, with strand- and coil-associated turns exhibiting higher dynamic propensity than helix-associated turns. Together, these results reveal how sequence composition and structural context jointly shape the conformational landscape of *β*-turns.

## Introduction

*β*-Turns are among the most common non-regular secondary structure elements in globular proteins, accounting for approximately 20−25% of residues and playing a central role in chain reversal, loop formation, and compact folding topologies [1–3]. Structurally, *β*-turns are defined by a four-residue segment (*i*−*i* + 3) in which the C*α* atoms of residues *i* and *i* + 3 are separated by less than 7.5 Å[2], often stabilized by a hydrogen bond between the backbone carbonyl of residue *i* and the amide of residue *i*+3. Early stereochemical analyses classified *β*-turns into a small number of canonical types (`I, II, I′, II′`), based on characteristic *ϕ*/*ψ* dihedral angle combinations of the central residues *i* + 1 and *i* + 2, establishing a geometric framework that remains widely used today[1, 4, 5].

Subsequent large-scale statistical analyses of high-resolution protein structures expanded this framework, identifying additional turn types (e.g., type VIII) and highlighting strong, position-specific amino-acid preferences within turns. In particular, glycine and proline were shown to play dominant but distinct roles: glycine frequently occupies the *i* + 2 position in turn types requiring access to disallowed regions of Ramachandran space, while proline is enriched at the *i* + 1 position where *ϕ* angle restriction stabilizes specific conformations [3, 5]. These studies established *β*-turns as sequence-encoded structural motifs, but largely treated them as static entities sampled from crystallographic ensembles.

More recent work has revisited *β*-turn classification using high-resolution structural datasets and clustering approaches in dihedral-angle space, revealing a richer landscape of turn conformations than captured by the classical scheme and emphasizing the heterogeneity of the traditionally defined “`type IV`” category [6]. In parallel, bioinformatics resources such as MapTurns have enabled systematic mapping of turn geometry and sequence context across modern protein structure databases, reinforcing the view that *β*-turns occupy a continuous conformational space rather than discrete, isolated states [7].

Despite these advances, *β*-turns are still most often described in terms of static frequencies and sequence propensities, while their dynamic behavior and interconversion pathways remain comparatively unexplored. Molecular dynamics simulations provide a natural framework for addressing this gap, allowing direct observation of transitions between turn types and enabling the dissection of how local sequence context modulates both turn stability and transition kinetics. Treating *β*-turns as dynamically interconverting states rather than fixed structural motifs offers a route toward understanding how sequence encodes not only preferred conformations but also the kinetic accessibility of alternative backbone geometries, with implications for folding, loop remodeling, and conformational switching in proteins.

## Results

### Clustering *β*-turns on the Cross Bond Ramachandran Plot reveals a hybrid cluster between turn type I and I′

To set the framework for our study we identified, in PDB structures [8], *β*-turns obeying the DSSP [9] pattern (`−|H|E, T, T, −|H|E`) and then applied K-means clustering [10] across the four dihedral angles of the central two residues in the segments. An optimal cluster number of six was identified by cross validation (Figs. S2 and S3), classifying turns into regions defined with the centers shown in Table 1.

**Table 1:**
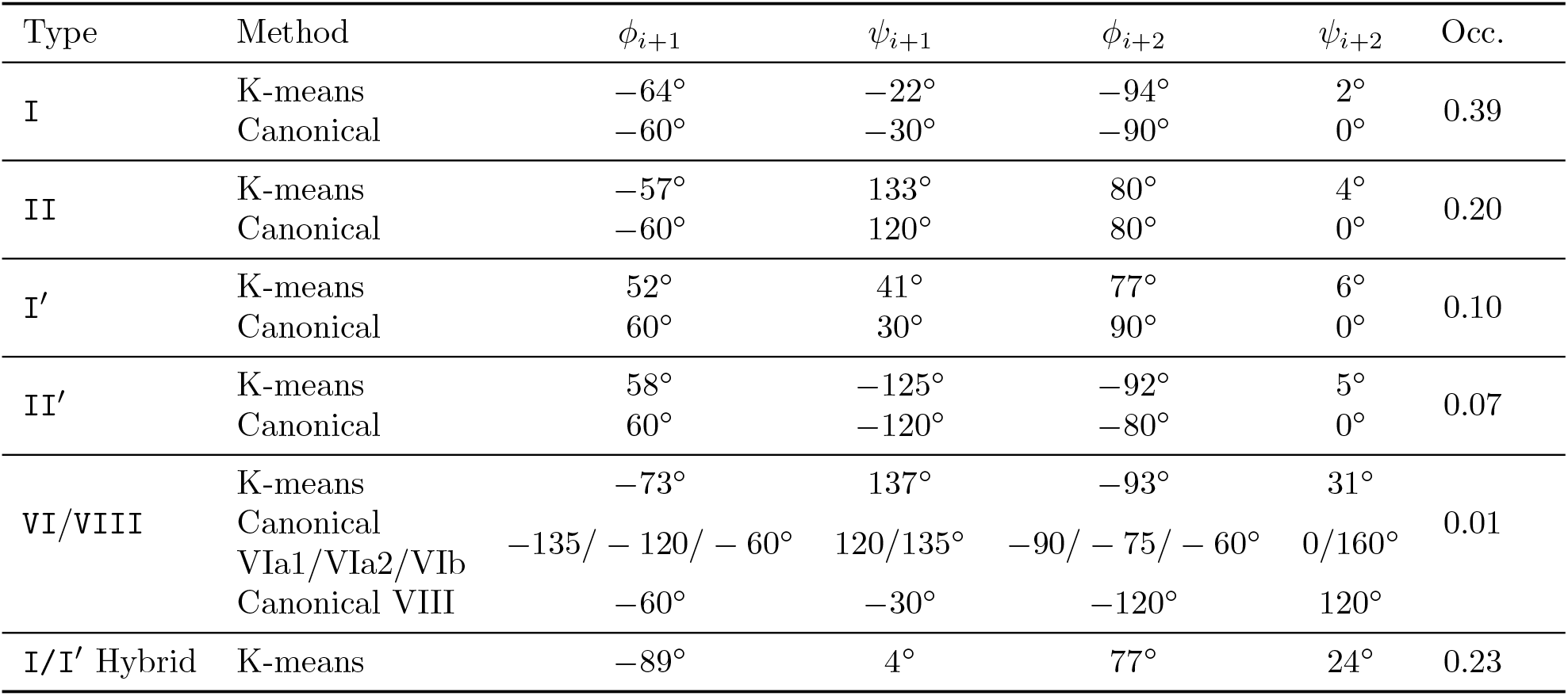
K-means centroids of six *β*-turn types aligned to canonical definitions.

While recent high-resolution analyses have proposed a finer subdivision of *β*-turn conformational space into a larger number of geometric subtypes (e.g., 18 clusters derived from ultra-high-resolution structures [6]), our goal here is different. Rather than maximizing structural subdivision in static datasets, we seek a compact representation that captures the dominant conformational basins observed in structural ensembles and molecular-dynamics trajectories. In this context, the six clusters can be interpreted as a coarse-grained description of *β*-turn conformational space that preserves the major canonical families while grouping closely related subtypes into kinetically and structurally meaningful regions.

These turn types clearly agree with the well known type `I, I′, II, II′` and additionally define a broad region of the Ramachandran Plot (Figs. 1 C-E and 2B, D, F, H) containing both cis and trans turns (Figs. 3D, S1) corresponding to the classically defined `VI` (cis) and `VIII` (trans) *β*-turn types. The characteristic geometries of these clusters are further illustrated by representative backbone conformations extracted from PDB structures (Fig. 1, Table S1)

**Figure 1:**
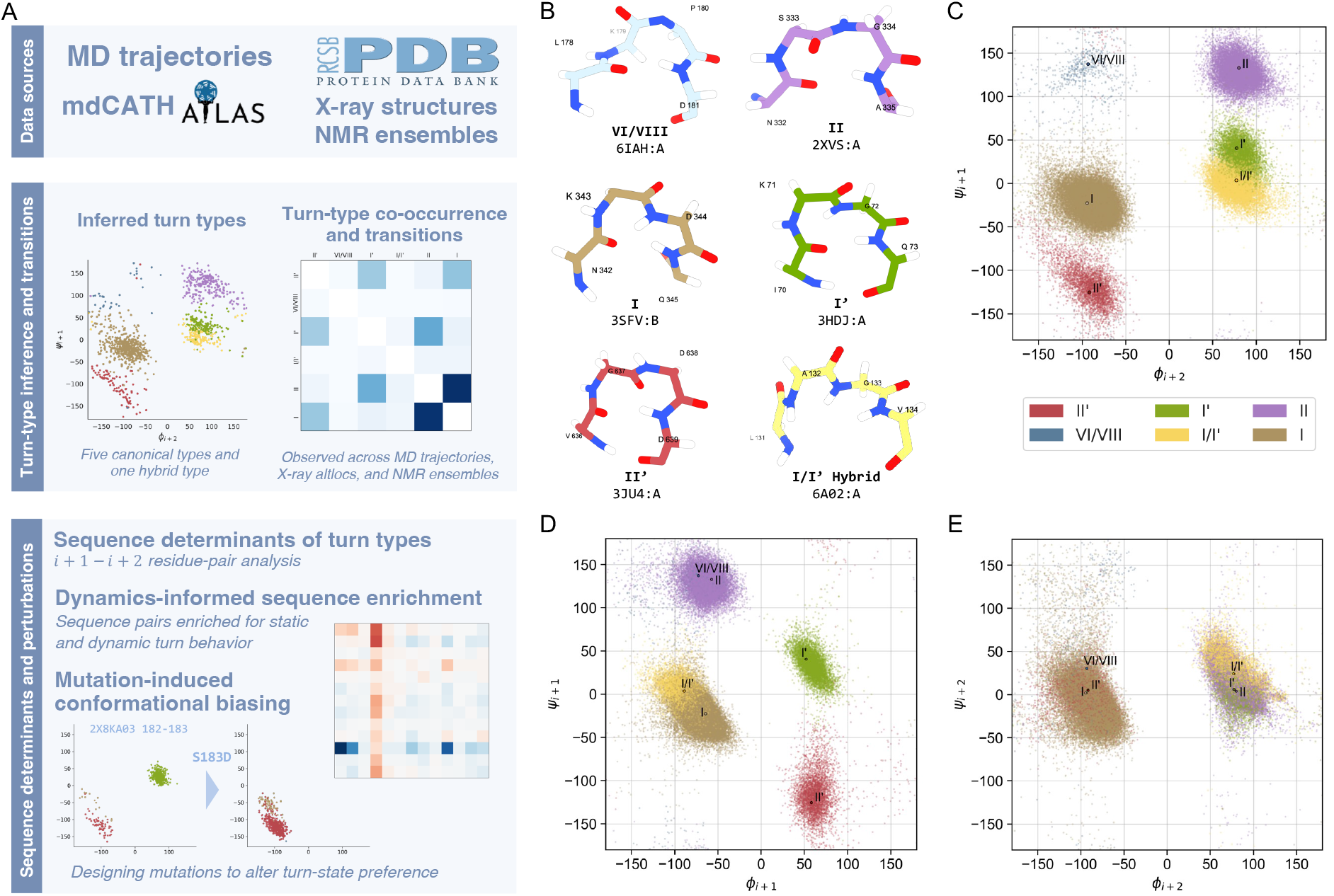
Data-driven classification of *β*-turn types. **(A)** Schematic overview of the study. The analysis integrates molecular dynamics simulations and experimental structures for *β*-turn classification and correlation analysis, and sequence preference characterization with mutation-based validation. *β*-turns were identified in PDB structures using a DSSP-based secondary-structure motif and classified by K-means clustering in dihedral-angle space. **(B)** Representative backbone conformations for each *β*-turn type extracted from PDB structures. Residues are colored according to turn type: `VI/VIII` (blue; `6IAH`, residues 178−181); `I` (beige; `3SFV`, 342−345); `II′` (red; `3JU4`, 636−639); `II` (purple; `2XVS`, 332−335); `I′` (green; `3HDJ`, 70−73); hybrid `I/I′` (yellow; `6A02`, 131−134). **(C-E)** Ramachandran plots show the starting conformation of *β*-turns from the first frame of the mdCATH dataset in dihedral-angle. *(C)* Cross-bond Ramachandran plot (*ϕ*_*i*+1_ versus *ψ*_*i*+1_), more clearly separates the identified clusters. Points are colored according to turn type as defined by K-means clustering. *(D)* Traditional Ramachandran plot centered on residue *i* + 1 (*ϕ*_*i*+1_ versus *ψ*_*i*+1_), and *(E)* centered on residue *i* + 2 (*ϕ*_*i*+2_ versus *ψ*_*i*+2_). Color coding is consistent throughout figures: `II′` (red), `VI/VIII` (blue), I′ (green), `I/I′` (yellow), `II` (purple), and `I` (beige).

**Figure 2:**
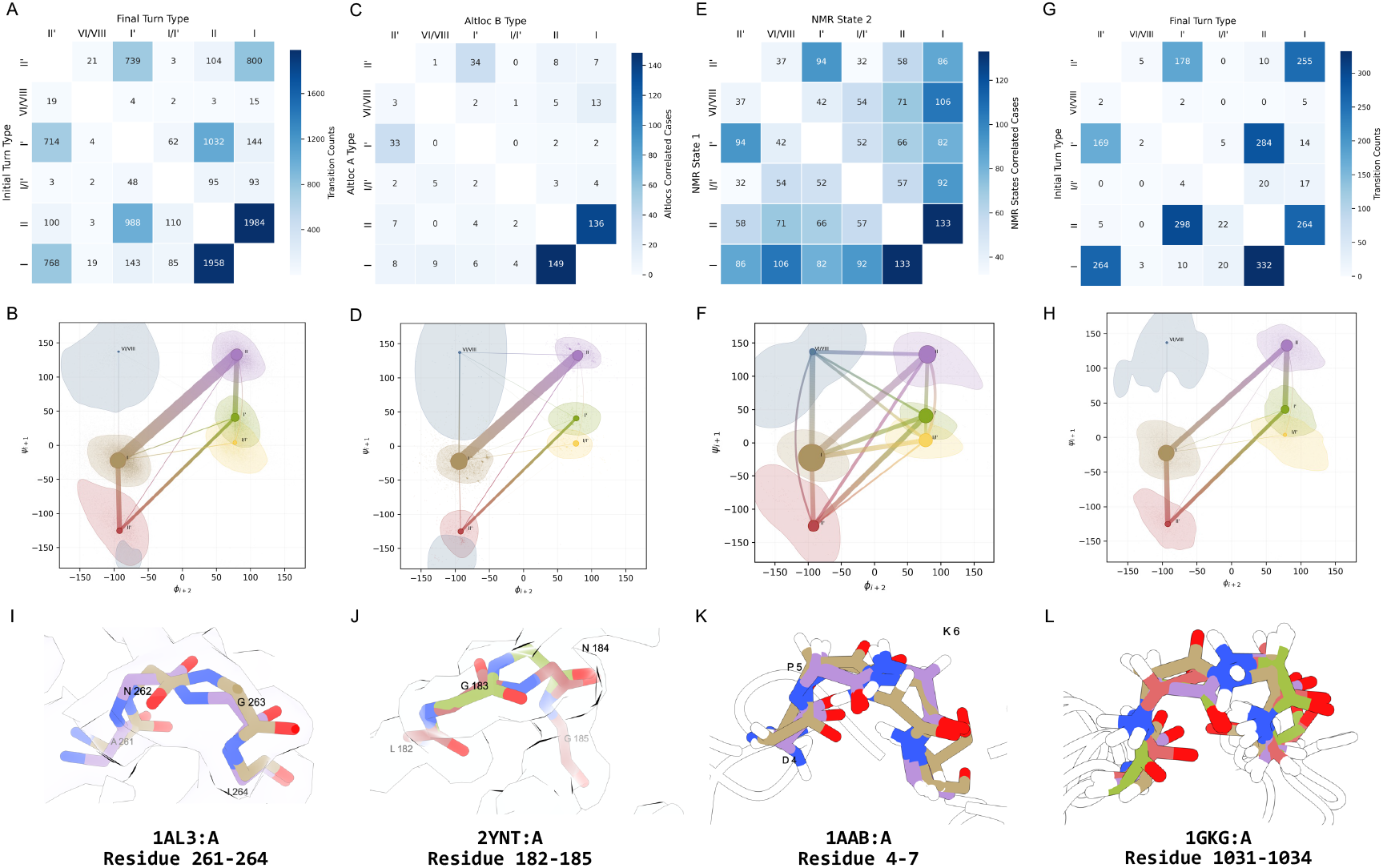
Patterns in *β*-turn dynamics observed in molecular dynamics match dual conformations observed in experimental structures. Turn-type transition counts were quantified across all mdCATH (320K) and ATLAS (300K) trajectories. *β*-turns adopting multiple conformations were identified from X-ray crystal structures based on alternate locations (altlocs) and from NMR ensembles based on state-dependent assignments. **(A, C, E, G)** Heatmaps showing transition or correlation frequencies between *β*-turn types for molecular dynamics trajectories in mdCATH (A) and ATLAS (G), X-ray altlocs (C), and NMR ensembles (E). **(B, D, F, H)** Network representations of the corresponding matrices, where nodes represent turn types and edges indicate transitions or correlations; node size scales with turn-type population, and edge width reflects transition frequency. Turn types are outlined to indicate their distribution in cross-dihedral space. Representative examples of *β*-turns adopting alternative conformations in X-ray crystal structures **(I, J)**, and NMR ensembles **(K, L)**. (I) PDB `1AL3`, chain A, residues 261−264, adopting type `I` (beige) and type `II` (purple) conformations. (J) PDB `2YNT`, chain A, residues 182−185, adopting type `II′` (red) and type `I′` (green) conformations. (K) PDB `1AAB`, chain A, residues 4−7, adopting type `I` (beige) and type `II` (purple) conformations. (L) PDB `1GKG`, chain A, residues 1031−1034, adopting type `I` (beige), type `II` (purple), type `II′` (red) and type `I′` (green) conformations. Turn types are colored as in Fig. 1.

**Figure 3:**
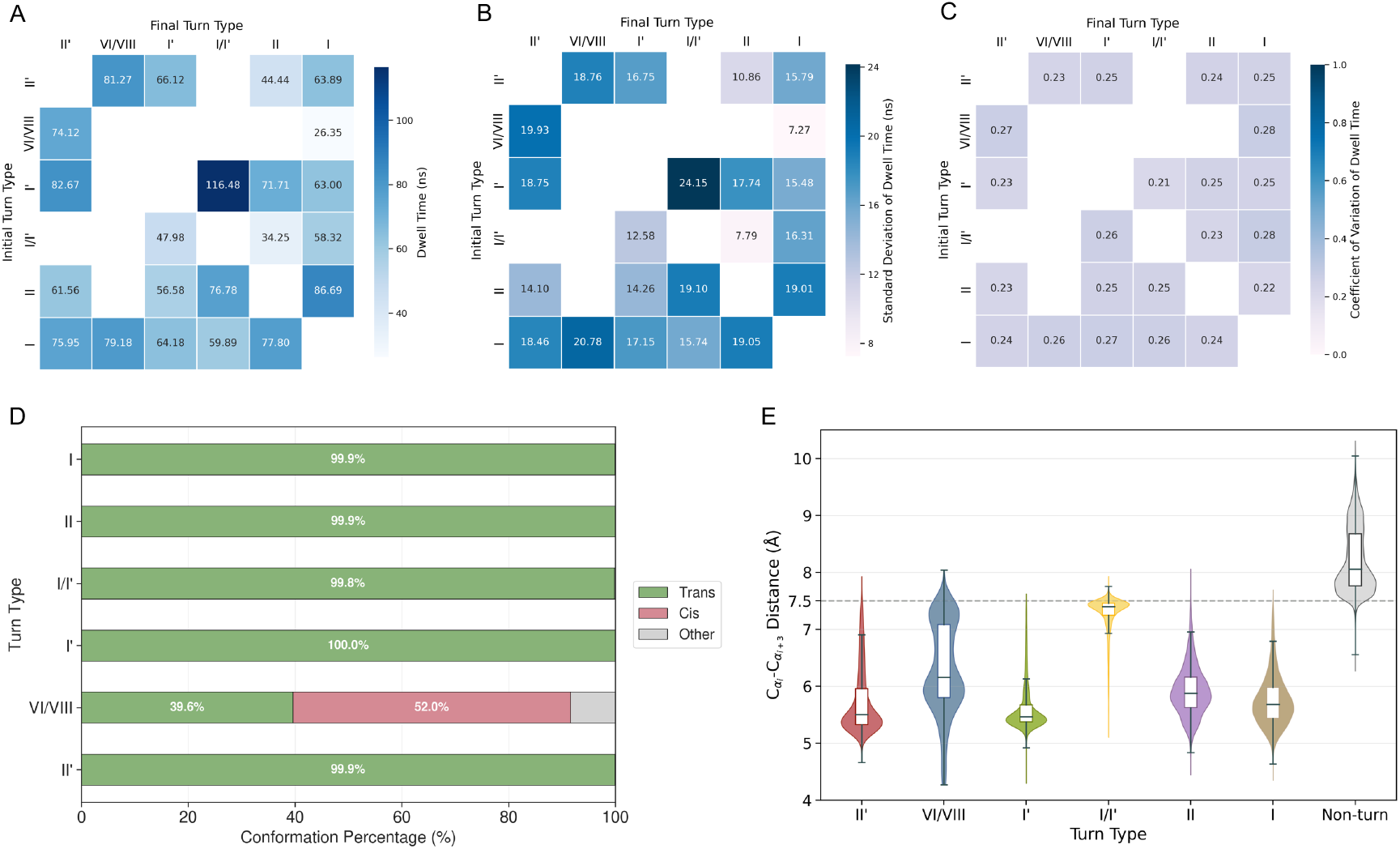
The hybrid I/I′ *β*-turn represents a transient conformational state. The `I/I′`turn type exhibits shorter dwell times and reduced structural compactness relative to other trans-conformation *β*-turn types (A–C). Average dwell time **(A)**, standard deviation **(B)**, and coefficient of variation **(C)** for transitions between turn types. Transitions out of the `I/I′` type occur on consistently shorter timescales than transitions into `I/I′` from other types, indicating reduced kinetic stability of the hybrid state. **(D)** cis–trans conformational composition of each turn type, evaluated using the peptide-bond dihedral angle *ω*_*i*+1_. The `VI` turn type contains a substantial fraction of cis conformations (52%), whereas all other turn types are exclusively trans. **(E)** Distribution of C*α*–C*α* distance between residues *i* and *i* + 3 computed over the trajectories and grouped by smoothed temporal turn-type labels. The `I/I′` type adopts a markedly larger average distance (7.3 Å) compared with other trans-conformation turn types (`II′, I′, II`, and `I`; 5–6 Å), indicating a less compact turn geometry.

In classical *β*-turn nomenclature, conformations that do not fall within the canonical *ϕ*/*ψ* ranges of Types `I, I′, II, II′, VI`, or `VIII` are grouped into the heterogeneous Type IV category [2, 3] that previous works have further subclassified into distinct types [6, 11]. We label this additional turn type as a hybrid `I/I′`, because it clusters near type `I` in the traditional Ramachandran plot of residue *i* + 1 and near type `I′` in both the cross-bond Ramachandran plot and the traditional Ramachandran plot of residue *i* + 2. Our recently described cross bond Ramachandran plot [12],centres on the apex of the *β*-turn and best separates the distinct turn types further justifying the usefulness of this representation (Fig. 1C). All further Ramachandran plots shown here use this cross bond representation.

### Dual conformation turns in experimental structures correspond to dominant interconversions between *β*-turn types observed in molecular dynamics

The central aim of this study is to investigate the nature of *β*-turn dynamics by analyzing transitions between the different *β*-turn clusters as observed in molecular dynamics (MD) trajectories. As described in the methods, we used mdCATH [13], a large scale database recently compiled to aid data driven computational biophysics and containing trajectories for 5, 398 protein domains, simulated in five replicates each at five temperatures from 320K to 450K, and complemented this analysis with simulations from the ATLAS dataset at 300K. Together, these datasets provide a comprehensive view of *β*-turn dynamics across diverse protein systems.

As a validation of the relevance of MD for exploring interchangeable *β*-turn conformations, we compared MD-derived transitions with alternative conformations observed in experimental structures. X-ray crystallography is intrinsically an ensemble measurement where the electron density captures the conformation modes arising from a lattice of billions of copies of the molecule being imaged. Regions of the protein chain occupying or dynamically interconverting between two discrete conformations can be observed in the electron density map as two distinct conformations that are modeled as alternately located atoms [14–16]. In parallel, NMR ensembles provide multiple structural models that capture conformational heterogeneity across solution states. We observe consistent patterns of interconversion across all datasets (Fig. 2). Molecular dynamics simulations from both mdCATH and ATLAS show that transitions are predominantly concentrated between specific turn-type pairs, most notably between types `I` and `II` and between `I′` and `II′`. These dominant transition pathways closely match the correlations observed between alternative conformations in X-ray structures. NMR ensembles display broader but qualitatively similar patterns of interconversion, with transitions spanning the same principal turn-type pairs. Together, these results demonstrate that the dominant *β*-turn transitions observed in MD simulations correspond closely to experimentally observed conformational heterogeneity, supporting the relevance of MD-derived transition networks for describing the conformational landscape of *β*-turns. Across datasets, transitions or correlations are predominantly observed between specific turn-type pairs. Molecular dynamics simulations show frequent transitions between `I-II, II-I′, I-II′`, and `I′-II′`, which are consistent with dominant correlations observed in X-ray structures. NMR ensembles show broader but similar patterns of interconversion.

### The hybrid I/I′ type *β*-turn represents a transient conformational state

We next examined whether the hybrid `I/I′` cluster identified in dihedral space exhibits distinct kinetic and structural properties relative to canonical *β*-turn types. Analysis of turns’ behavior across trajectories revealed that the `I/I′` type displays markedly shorter dwell times compared with other trans-conformation turn types (Fig. 3A). Directional transition comparison further demonstrates that transitions out of the `I/I′` state occur on consistently shorter timescales thantransitions into `I/I′` from other turn types. This asymmetry indicates reduced kinetic stability and suggests that `I/I′` functions as a transient intermediate along conformational exchange pathways rather than a long-lived structural basin. Importantly, these transition patterns, particularly those involving the `I/I′` state, are preserved across a range of temporal smoothing windows (Fig. S1C), indicating that the transient behavior of `I/I′` reflects a reproducible intermediate rather than noise arising from short-lived fluctuations. To exclude the possibility that the observed instability arises from cis-trans heterogeneity, we quantified peptide-bond geometry using the *ω*_*i*+1_ dihedral angle (Fig. 3D). As expected, type `VI/VIII` contains a substantial fraction of cis conformations (52%), whereas `I/I′` and all other non-`VI` types are exclusively trans. Thus, the transient behavior of `I/I′` cannot be attributed to cis-trans isomerization. Structural compactness analysis further distinguishes the hybrid state. The C*α*-C*α* distance between residues *i* and *i* + 3 is markedly larger for `I/I′` (mean ≈ 7.3 Å) than for other trans-conformation turn types (5−6 Å; Fig. 3E), indicating a less compact backbone geometry. Together, these kinetic and geometric features support a model in which the `I/I′` hybrid cluster represents a shallow, weakly stabilized conformational state that facilitates interconversion between neighboring *β*-turn geometries.

### Analyzing amino acid preferences in static and dynamic turns

Every *β*-turn in the mdCATH dataset was labeled as the most frequently adopted conformation type across the trajectory. We analysed the sequence features characteristics of each turn type following this labeling and found overall consistency with previously described *β*-turn type specific amino acid propensities [2, 3, 17]. Fig. 4 shows that type I turns stand out as, representing the largest population, having broad sequence tolerance. Type `I′`, `II` and `I/I′` turns share a strong enrichment of glycine at position *i* + 2, while they exhibit distinct characteristics on position *i* + 1. Type `I′` is enriched with `ASN` and `ASP` at position *i* + 1, type `II` preference `PRO` and `GLU` and type `I/I′` shows enrichment for `ALA` at position *i* + 1. Type `II′` uniquely favors Glycine at position *i* + 1, with GLY-ASP, GLY-ASN, and `GLY-SER` enrichment. Type `VI/VIII` turns are distinguished by a pronounced preference for `PRO` at *i* + 2, consistent with the known association of type `VI` turns (including `VIa1, VIa2`, and `VIb`) with cis-peptide configurations. It is noticeable in Fig. 6 that turn pairs identified as having greater propensity to dynamically interconvert demonstrate complementary sequence requirements in the sense that the most prominent enrichment signal for type `I` and type `II′` are at position *i*+1 whilst the most prominent enrichment signal for the dynamic counterpart, type `II` and type `I′`, respectively, are at the adjacent position *i* + 2. This potentially creates a setting where a particular turn can comply with the strongest sequence constraints for both turn types. It is also worth noting that for turn types `I′` and `II′` numerous weaker enrichment signals are correlated, most noticeably for combinations of `ASN, ASP` and `GLU` at positions *i* + 1 and *i* + 2.

**Figure 4:**
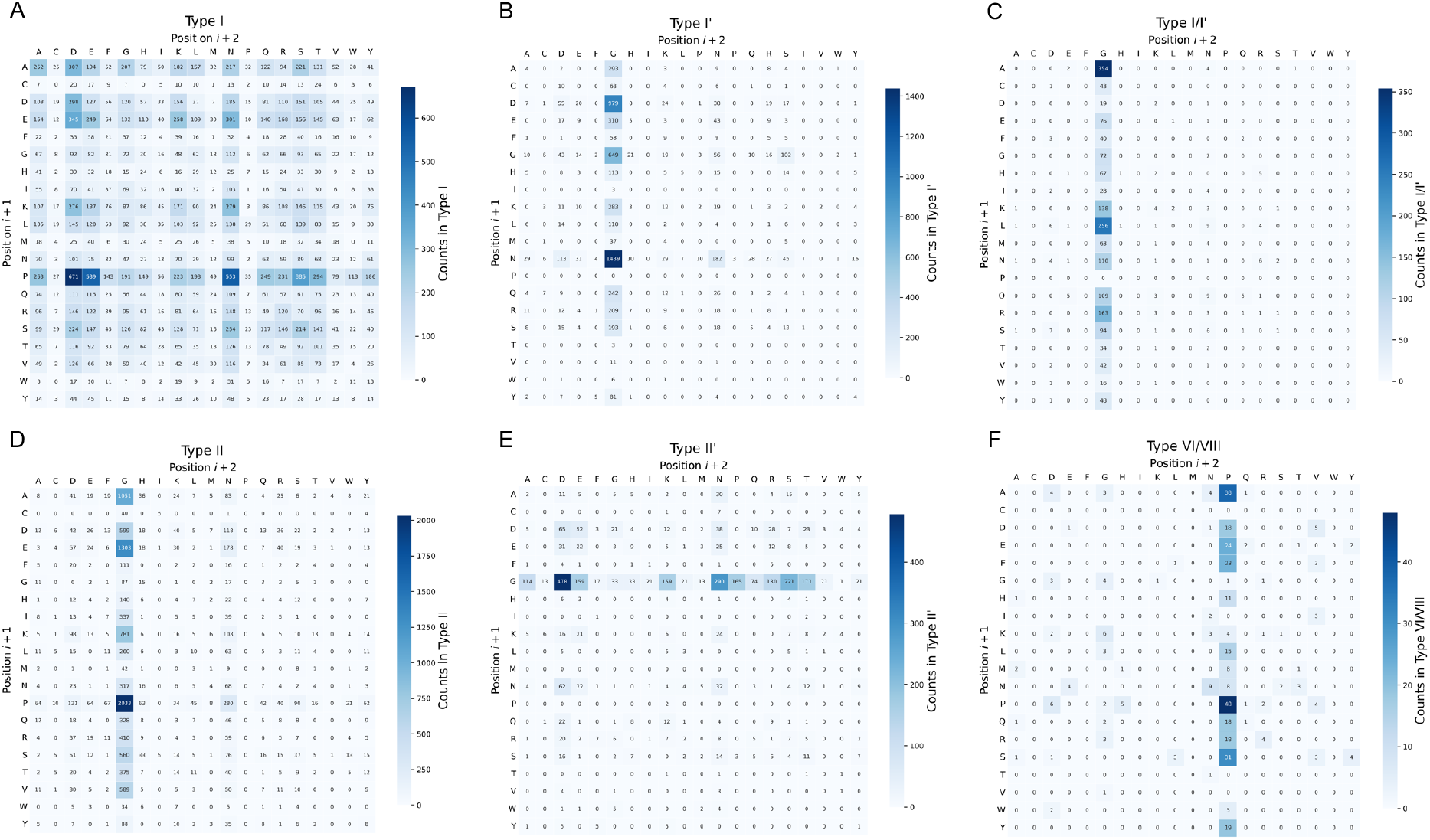
Pairwise amino acid composition of *β*-turn types. Matrices show the occurrence counts of amino acid pairs at positions *i* + 1 and *i* + 2 for each *β*-turn type. Each cell represents the number of turns containing a specific residue pair within that type. Values are colored using a blue gradient color, where lighter colors indicate low counts and darker colors indicate higher counts. Distinct *β*-turn types exhibit characteristic amino acid preferences at positions *i* + 1 and *i* + 2. **(A)** Type `I` turns display broad amino acid preferences, with `PRO, ALA, LYS, ASP, GLU`, and `SER` enriched at position *i* + 1 and `ASP` most frequent at position *i* + 2, followed by `ASN, GLU`, and `SER`. **(B)** Type `I′` turns are enriched for Glycine at position *i* + 2, with `ASN, ASP`, and `GLY`, and most frequently observed at position *i* + 1. **(C)** The hybrid `I/I′` type shows an almost exclusive presence of Glycine at position *i* + 2, and enriched for `ALA, LEU, ARG` at position *i* + 1. **(D)** Type `II` turns preferentially contain `GLY` at position *i* + 2, followed by `ASN`; position *i* + 1 is enriched for PRO, followed by `ALA, GLU`, and `LYS`. (E) Type `II′` turns show a high frequency of `GLY` at position *i* + 1 and enrichment of `ASP, ASN, SER, GLU` and `PRO` at position *i* + 2. (F) Type `VI/VIII` turns show a strong preference for `PRO` at position *i* + 2, while position *i* + 1 is enriched for `PRO, ALA, SER, GLU`, and `PHE`.

To further investigate how amino acid sequence influences conformational dynamics, turns were first classified based on transition counts between turn type conformations along trajectories: segments exhibiting no transitions were defined as static, whereas those making at least three transitions were defined as dynamic. Table 2 summarizes the distribution of static and dynamic turns within each type. The hybrid `I/I′` type exhibits the strongest dynamic tendency, with more than half of the turns of this type classified as dynamic, consistent with its transient conformational character. In contrast, type `VI/VIII` turns are predominantly static.

**Table 2:** Distribution of static and dynamic *β*-turn segments across different turn types.

| Type | Size | Static |  | Dynamic |  |
| --- | --- | --- | --- | --- | --- |
|  |  | Count | % | Count | % |
| II' | 3234 | 1458 | 45.1% | 1161 | 35.9% |
| VI/VIII | 417 | 315 | 75.5% | 58 | 13.9% |
| I' | 6741 | 4159 | 61.7% | 1597 | 23.7% |
| I/I' Hybrid | 1920 | 421 | 21.9% | 1221 | 63.1% |
| II | 13575 | 8652 | 63.7% | 2908 | 21.4% |
| I | 25933 | 18100 | 69.8% | 4278 | 16.5% |

To determine whether particular residue pairs bias a turn towards dynamic or static behavior, we compared the composition of amino acid pairs across static and dynamic segments (Fig. S4), and calculated enrichment scores, i.e. the difference between the relative frequency of each amino acid pair in dynamic versus static turns (Fig. 5). Notably, residue pairs that are highly common within a given turn type do not contribute equally to conformational flexibility. Within the same structural class, certain frequent pairs are biased toward dynamic behavior, whereas others preferentially stabilize static conformations. For example, in type `II′` turns, the `GLY-SER` pair exhibits strong dynamic enrichment, whereas GLY-ASP, GLY-ASN, and `GLY-PRO` are preferentially associated with static behavior. A similar within-type contrast is observed for type `I′` turns: although `ASN-GLY` and `GLY-GLY` are common in this class, they are enriched in static segments, whereas `GLY-SER` correlates with increased dynamic behavior. Such correlation indicates that turns with `GLY-SER` pairs are likely to be associated with transition between type `I′` and `II′`. In type `II` turns, the most populated `PRO-GLY` pair strongly favors static conformations, while pairs such as `ALA-GLY,GLU-GLY`, and `SER-GLY` are enriched in dynamic segments. In type `I` turns, proline at position *i* + 1 correlates with static behavior, whereas Glycine containing pairs at *i* + 2 are associated with increased conformational exchange.

**Figure 5:**
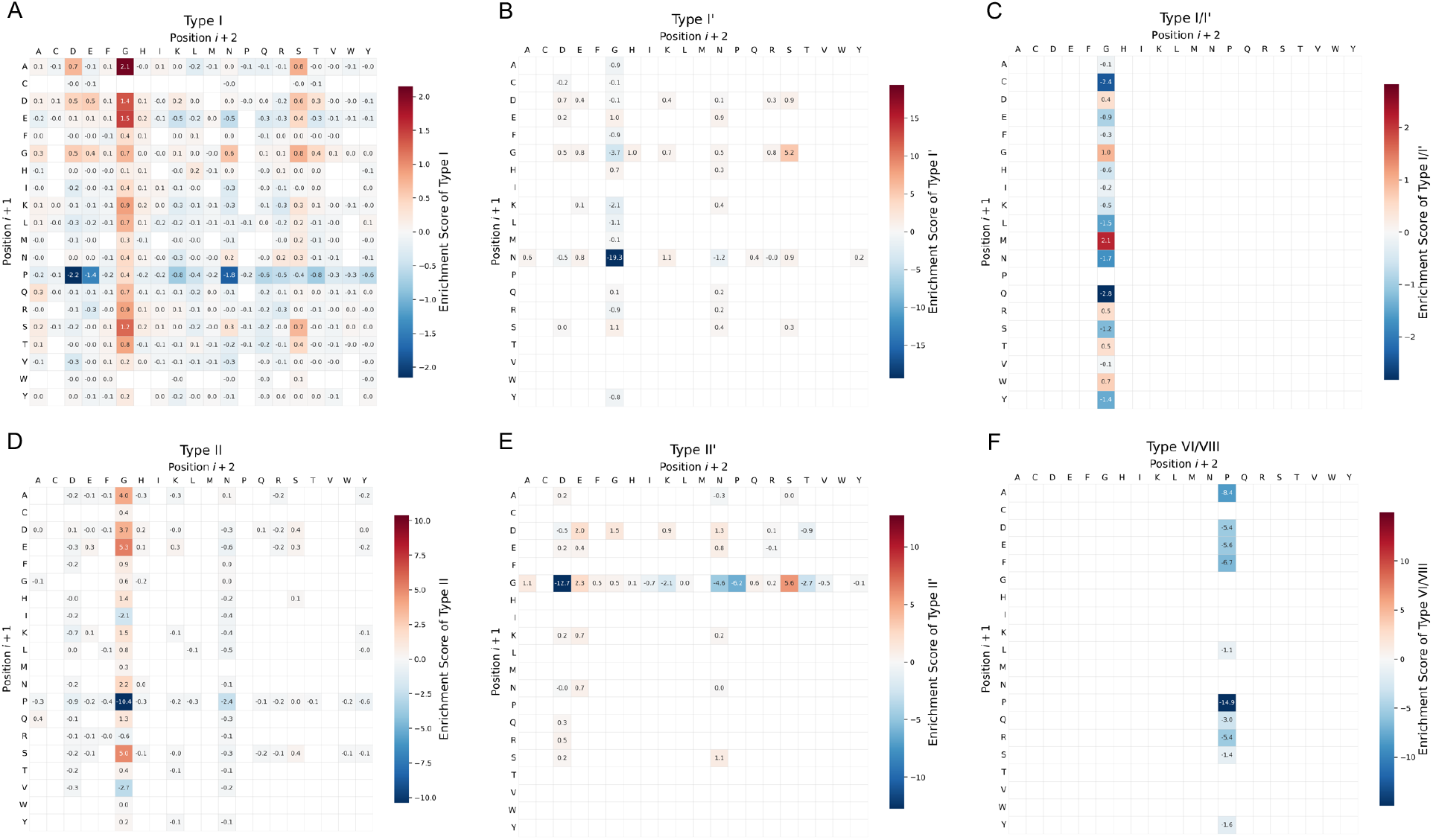
Dynamic and static biases within turn-type-specific amino acid preferences. Enrichment scores are shown for each *i* + 1 and *i* + 2 pair within each turn type. Enrichment scores were calculated as the percentage of each amino acid pair observed in dynamic turns minus that in static turns and shown in a blue to red color gradient. Positive values indicate enrichment in dynamic turns (red) and negative values indicate enrichment in static turns (blue). Scores for residue pairs with a total occurrence count of fewer than 10 (dynamic and static combined) were not displayed to avoid over interpretation of sparse data.

Notably, although type `VI/VIII` turns are globally static, their dominant residue pairs (`PRO-PRO` and `ALA-PRO`) reinforce this stability, and dynamic instances are too rare to define robust sequence determinants. The hybrid `I/I′` type, despite being globally dynamic, also shows internal sequence-dependent modulation: `ALA-GLY` and `MET-GLY` pairs are preferentially associated with dynamic turns, whereas `GLN-GLY` correlates with static behavior. Together, these results demonstrate that dynamic behavior is not determined solely by turn type but is modulated by specific residue pairs within each structural class.

To more directly test whether residue-pair composition causally influences *β*-turn dynamics, at least in silico, we introduced targeted substitutions that convert dynamic-enriched pairs to static-enriched pairs and vice versa (Fig. 6 and Table S3) and reran MD simulations. Conformational distributions were evaluated using the cross-bond Ramachandran representation together with type assignments across time.

**Figure 6:**
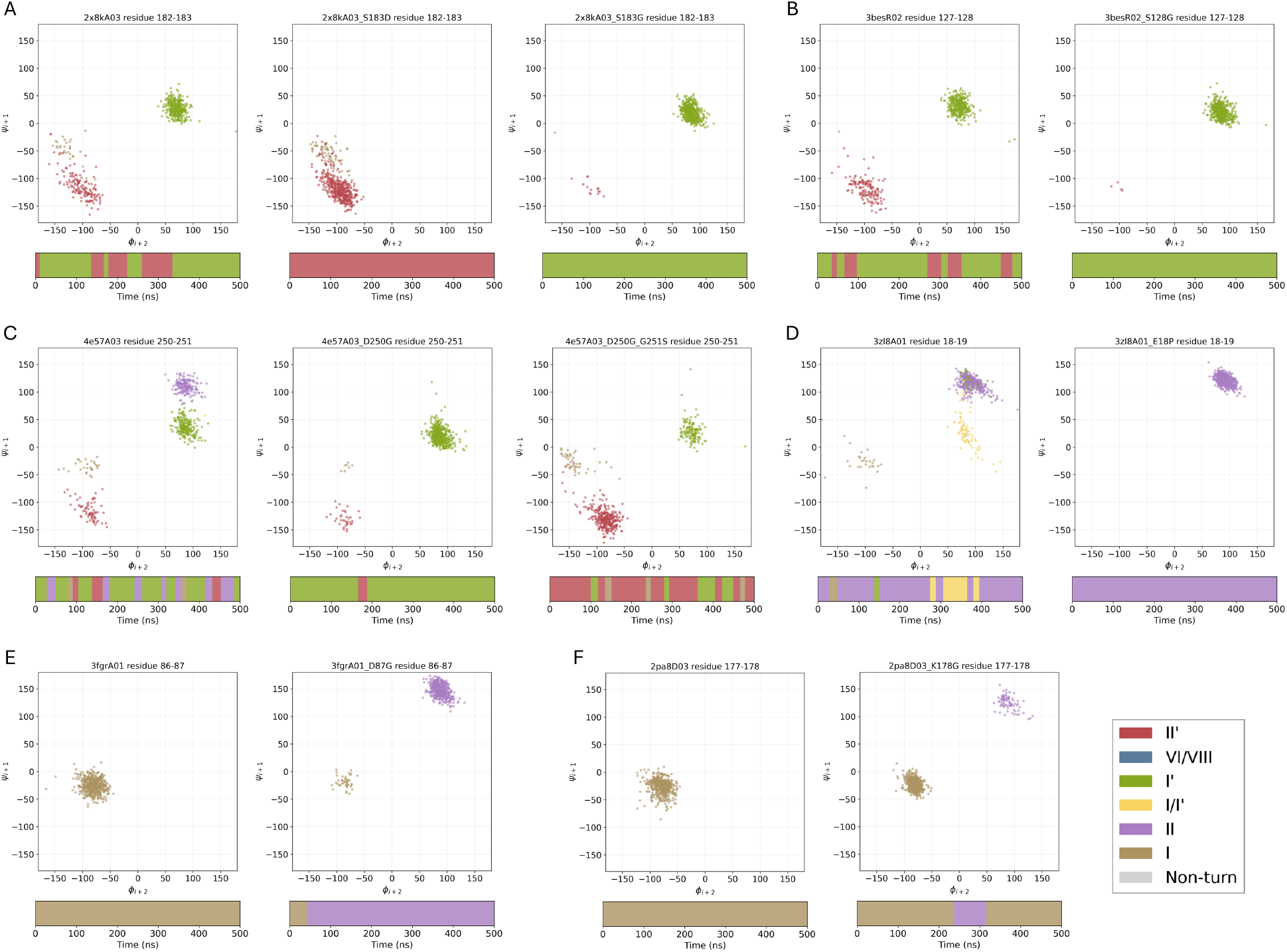
Targeted amino acid substitutions alter *β*-turn dynamics. Mutations converting dynamic-enriched residue pairs to static-enriched pairs (A-D) restrict conformational heterogeneity, whereas substitutions converting static-enriched pairs to dynamic-enriched pairs (C, E, F) increase turn-state plasticity. The top plots show the distribution of *ϕ*_*i*+2_ versus *ψ*_*i*+1_ at 500 frames in the cross bond Ramachandran plot, and the bottom plots show the turn type assignments across time. **(A)** CATH `2X8KA03` (residues 182− 183). The wild-type `GLY-SER` pair adopts both type `I′` and `II′` conformations (left). The S183D substitution (`GLY-ASP`) restricts the turn to type `II′` (middle), whereas S183G (`GLY-GLY`) stabilizes the type `I′` conformation (right). **(B)** CATH `3BESR02` (residues 127 −128). The wild-type `GLY-SER` pair transitions between type `I′` and `II′` states (left). The S128G substitution (`GLY-GLY`) confines the turn to the type `I′` conformation (right). **(C)** CATH `4E57A03` (residues 250-251). The wild-type (`ASP-GLY`) adopts type `I, II, I′, II′` conformations (left), D250G (`GLY-GLY`) mutation restricts the turn to mainly type `I′` (middle), and D250G-G251S (`GLY-SER`) mutation allows transitions between `I′, II′` and `I` (right). **(D)** CATH `3ZL8A01` (residues 18− 19). The wild-type `GLU-GLY` pair adopts type `I, II`, and hybrid `I/I′` conformations (left). Substitutions E18P (`PRO-GLY`, right) restricts the turn to type `II`. **(E)** CATH `3FGRA01` (residues 86−87). The wild-type `PRO-ASP` pair adopts a type `I` conformation (left). The D87G substitution (`PRO-GLY`; right) enables both type `I` and `II` states. **(F)** CATH `2PA8D03` (residues 177−178). The wild-type `GLU-LYS` pair adopts type `I` only (left). The K178G substitution (`GLU-GLY`) enables both type `I` and `II` conformations (right).

Consistent with the enrichment analysis (Fig. 5), substitutions that replaced dynamic-enriched pairs with static-enriched pairs markedly reduced conformational heterogeneity (Fig. 6A–D). In mdCATH structures `2X8KA03` and `3BESR02`, wild-type `GLY-SER` pairs sampled both type `I′` and `II′` states, whereas substitution to `GLY-ASP` or `GLY-GLY` restricted the turns to a single dominant conformation. Similarly for `3ZL8A01`, the wild-type `GLU-GLY` pair accessed type `I, II`, and hybrid `I/I′` conformations, while replacement with `VAL–GLY` or `PRO–GLY` confined the turn to type `II`. Likewise, in `4E57A03` the wild-type `ASP-GLY` pair samples multiple turn types (`I, II, I′`, and `II′`),while mutation to `GLY-GLY` confines the turn primarily to type `I′`. Notably, further substitution to `GLY-SER` reintroduces conformational transitions, allowing the turn to sample multiple states again.Conversely, substitutions converting static-enriched pairs to dynamic-enriched pairs expanded conformational sampling (Fig. 6E, F). In `3FGRA03`, replacement of `PRO-ASP` with `PRO-GLY` enabled coexistence of type `I` and `II` states. Likewise, in `2PA8D03`, substitution of `GLU-LYS` with `GLU-GLY` introduced transitions between type `I` and `II` states.

Across all systems examined, the direction of change in conformational plasticity was consistent with the enrichment biases identified within each turn type. These results suggest that residue-pair composition is not merely correlated with *β*-turn dynamics but can actively modulate the conformational landscape in a predictable manner.

### Sequence design models capture conformational bias in *β*-turns

To assess whether sequence preferences associated with alternative *β*-turn conformations are captured by structure-conditioned sequence models, we adopt a conformational bias framework [18]. Here, sequence likelihoods are compared across backbone conformations, and differences in model-assigned log-probabilities are interpreted as a measure of conformational bias. We apply this approach using ProteinMPNN [19], comparing to dominant (static) conformations with alternative structures generated by AlphaFold3 [15, 16].

ProteinMPNN-derived log-probability differences positively correlate with sequence prevalence observed in molecular dynamics simulations, as measured by the log_2_ prevalence ratio of residue pairs at positions *i* + 1 and *i* + 2 (Fig. 7A–D). This indicates that structure-conditioned models capture conformational preferences at the level of local *β*-turn motifs. However, dynamic- and static-enriched residue pairs do not show clear separation, suggesting that these scores do not fully capture sequence determinants of conformational dynamics.

**Figure 7:**
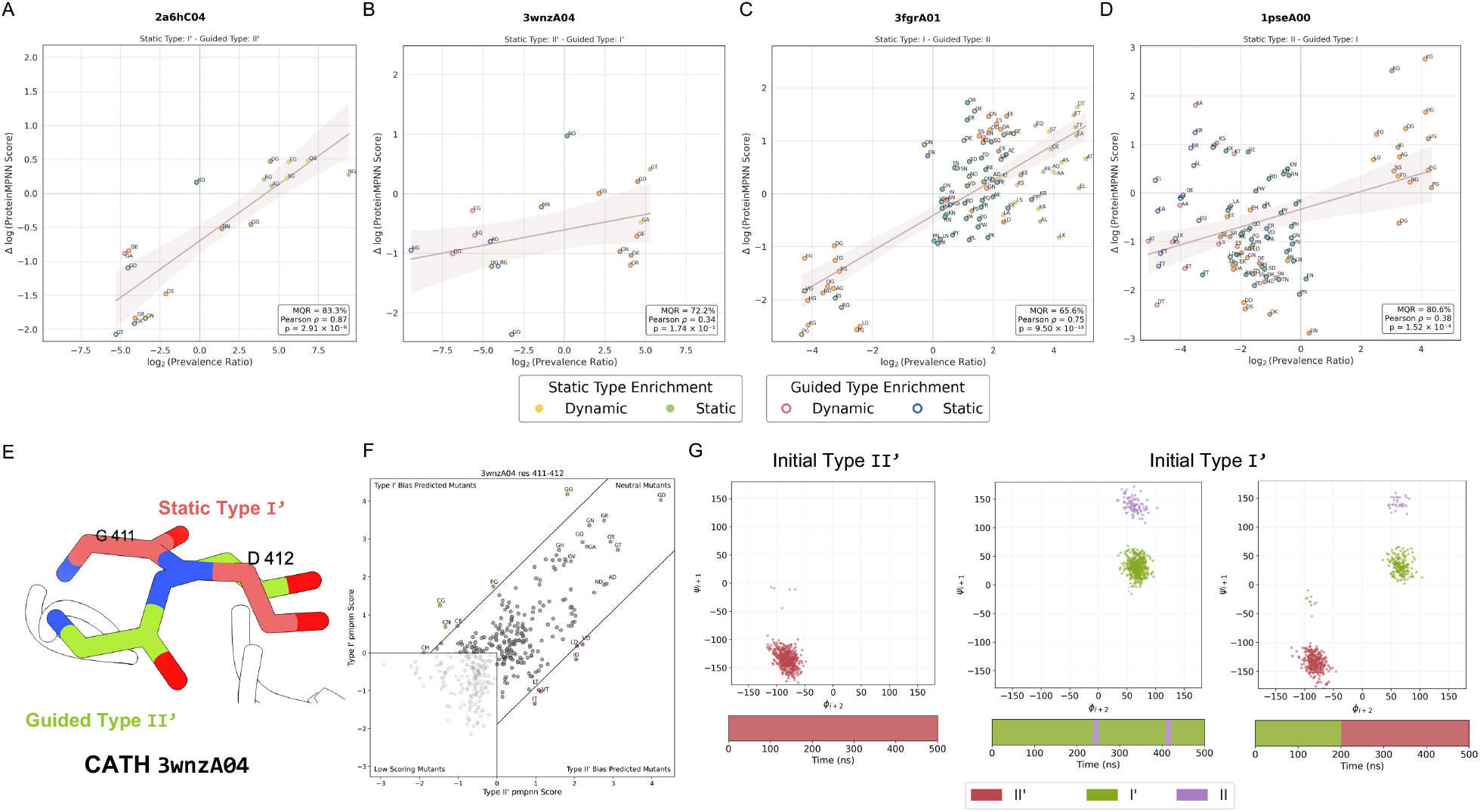
ProteinMPNN captures sequence preferences associated with *β*-turn conformations but only partially reflects dynamic behavior. **(A–D)** Correlation between ProteinMPNN-derived sequence preference differences and observed sequence prevalence across *β*-turn conformations. For each pair at positions *i*+1 and *i*+2, the log-probability difference between static and AF-guided conformations is plotted against the log_2_ prevalence ratio from mdCATH (Fig. 4). Pearson correlations and matched quadrant ratios (MQR; the fraction of points with consistent sign between the two axes) indicate that ProteinMPNN captures overall sequence preferences associated with alternative conformations. Points are colored by dynamic/static enrichment (Fig. 5): dynamic-enriched pairs (yellow fill for static, pink edges for guided) and static-enriched pairs (green fill for static, blue edges for guided). The lack of clear separation suggests that Protein-MPNN does not fully capture sequence determinants of conformational dynamics. **(E–G)** *In silico* mutation analysis for CATH `3WNZA04` (residues 410–413). **(E)** Backbone structures of residues 411–412 in the static (red; type `I′`) and AF-guided conformation (green; type `II′`). **(F)** Protein-MPNN conformational bias scores between `I′` and `II′`, showing similar high scores for the `GLY-ASP` (GD) pair. **(G)** Molecular dynamics simulations initialized from each conformation for the same sequence reveal distinct behaviors: trajectories starting from `II′` remain stable, while those from `I′` predominantly retain `I′` with occasional transitions toward `II` or `II′`.

Extending this analysis, we relate conformational bias to dynamic behavior. We hypothe-size residue pairs with strong bias tend to exhibit stable (static) conformations, whereas balanced scores correspond to increased conformational variability. This is illustrated by the `GLY-ASP` example (CATH `3WNZA04`), which shows similar ProteinMPNN scores for type `I′` and `II′` conformations (Fig. 7E–F) and exhibits distinct trajectories depending on initialization (Fig. 7G). However, not all predicted biases translate into stability some residue pairs favoring a conformation fail to maintain it in simulation (Table S2). These results indicate that sequence-conditioned likelihoods capture conformational preferences but do not fully determine dynamic behavior, which depends on additional sequence-structure interactions governing local energetics.

### Effects of context on dynamics: differences in transition across local secondary structures

To assess how the local structural context, adjacent to the turn, influences *β*-turn conformational exchange, we quantified transition counts between turn types across distinct secondary-structure environments defined by flanking DSSP states (Table 3). Turns were grouped according to the secondary-structure pattern surrounding the central `TT` segment, allowing comparison of transition behavior under helix-, strand-, or coil-adjacent contexts. Turns embedded within helical environments (`HTTH`) exhibit the strongest structural persistence, with more than 77% of segments remaining static and only ~ 12% classified as dynamic. Similarly, turns flanked by mixed helix– coil or strand–coil contexts (`HTTE/ETTH` and `-TTE/ETT-`) remain predominantly static. In contrast,strand-flanked turns (`ETTE`) and coil-flanked turns (`-TT-`) display markedly higher conformational variability, with around 27% fractions being dynamic.

**Table 3:** Composition of static and dynamic turns under different local secondary structures.

| Structure Context | Size | Static |  | Dynamic |  |
| --- | --- | --- | --- | --- | --- |
|  |  | Count | % | Count | % |
| HTTE/ETTH | 687 | 498 | 72.5% | 127 | 18.5% |
| HTTH | 1570 | 1209 | 77.0% | 188 | 12.0% |
| -TTE/ETT- | 5388 | 3670 | 68.1% | 945 | 17.5% |
| ETTE | 11896 | 6359 | 53.5% | 3634 | 30.6% |
| HTT-/-TTH | 21235 | 13450 | 63.3% | 5091 | 24.0% |
| -TT- | 25180 | 13784 | 54.7% | 6853 | 27.2% |

Transition behavior varies markedly with structural environment (Fig. 8). Turns embedded within helical contexts (`HTTH`) exhibit relatively low overall transition counts, with exchanges largely confined to type `I ↔ II′` and `I ↔ II` pathways. A similar restriction is observed for `ETTH` and `HTTE` motifs, suggesting that the presence of structured flanking elements constrains accessible conformational transitions.

**Figure 8:**
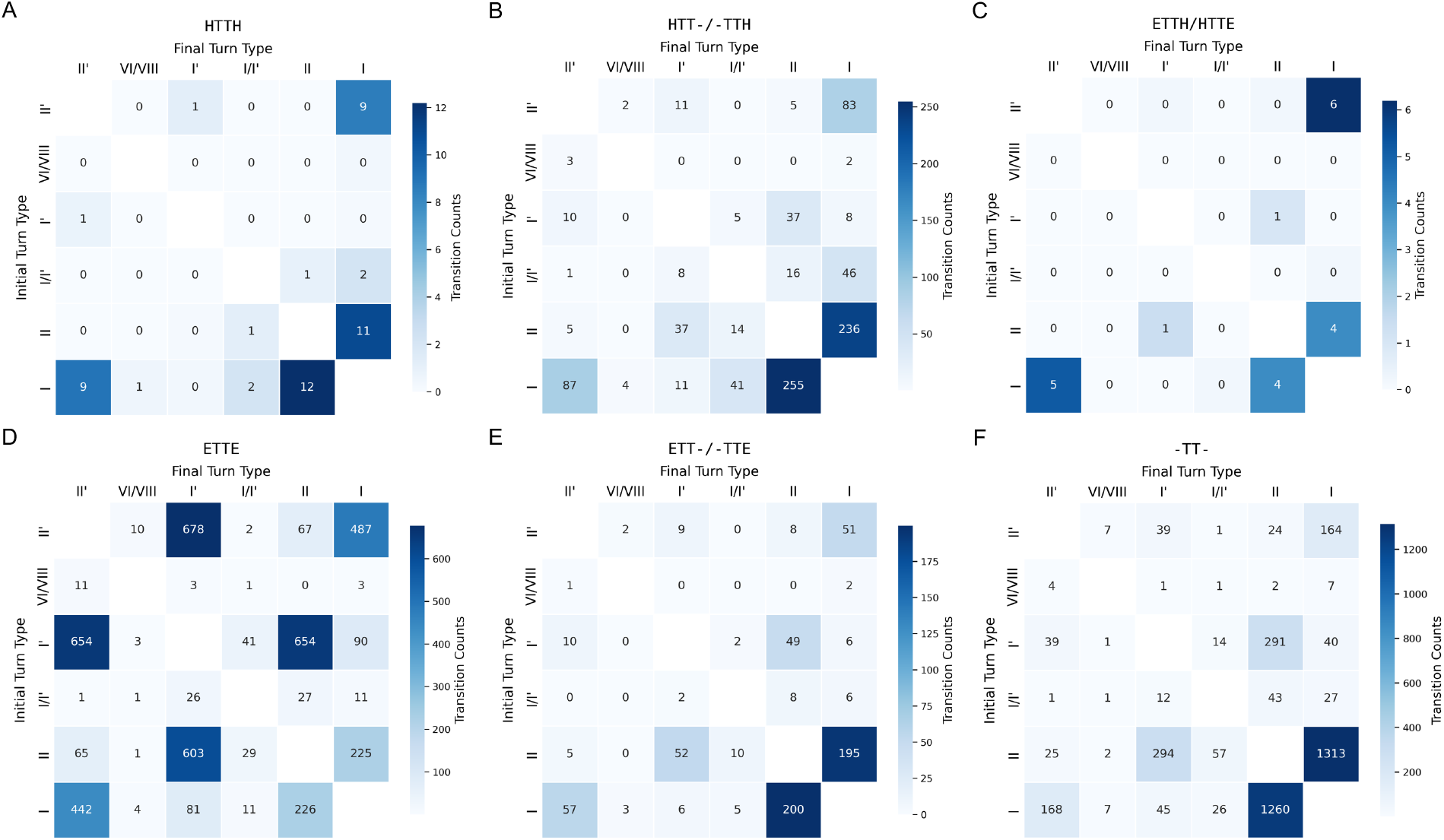
Transition patterns of *β*-turn types in different local secondary-structure contexts. Transition count matrices are shown for *β*-turn segments categorized by flanking secondary-structure motifs defined by DSSP. **(A)** `HTTH` turns exhibit overall low transition counts, primarily between type `I` and `II′` and between type `I` and `II`. **(B)** `HTT-` and `-TTH` turns show type `I ↔ II` transitions as the dominant exchange pathway. **(C)** `ETTH` and `HTTE` turns display relatively low transition counts, with transitions concentrated between type `I` and `II′` and between type `I` and `II`. **(D)** `ETTE` turns exhibit the broadest distribution of transitions, with frequent interconversion between type `I′` and `II′`. **(E)** `ETT-` and `-TTE` turns are dominated by `I ↔ II` transitions. **(F)** `-TT-`turns are similarly dominated by type `I ↔ II` transitions while exhibiting higher overall transition counts.

In contrast, turns situated between *β*-strands (`ETTE`) display the broadest transition spectrum, with frequent interconversion between type `I′` and `II′` states. This increased diversity suggests that strand–turn–strand architectures permit greater conformational plasticity within the turn segment.Intermediate contexts (HTT-, -TTH, ETT-, -TTE, and `-TT-`) are predominantly characterized by type `I ↔ II` transitions, indicating that these two geometries form a common exchange axis across partially structured or flexible environments.

Collectively, these results demonstrate that *β*-turn conformational dynamics are not governed solely by intrinsic residue-pair composition but are further modulated by the surrounding secondary-structure framework, which selectively constrains or facilitates specific transition pathways.

## Conclusion

*β*-turns have long been described as discrete structural motifs defined by characteristic backbone dihedral angles and sequence propensities. Our analysis of the large-scale molecular dynamics trajectories from mdCATH supports a more dynamic view. The importance of the dynamic view is highlighted by the identification of a hybrid `I/I′` cluster that behaves as a transient conformational intermediate. Although geometrically adjacent to canonical turn types, this state exhibits markedly shorter dwell times and reduced structural compactness, with an expanded C*α*–C*α* separation between residues *i* and *i* + 3. It appears to function as a kinetic bridge that facilitates exchange between neighboring conformations. Such intermediate states are difficult to recognize in static structural datasets but emerge clearly in time-resolved analyses.

More generally our results demonstrate that sequence encodes not only preferred *β*-turn conformations but also dynamic behavior. Within a given turn type, specific residue pairs at positions *i*+1 and *i* + 2 bias turns toward either static persistence or dynamic exchange. Targeted substitutions that convert dynamic-enriched pairs to static-enriched pairs reduce conformational heterogeneity in simulation, whereas the reverse substitutions expand accessible states. Although performed in silico, these perturbations consistently follow enrichment-derived predictions, suggesting that residue-pair composition can tune backbone plasticity in a predictable manner. Consistent with this view, structure-conditioned sequence models such as ProteinMPNN capture conformational preferences between different *β*-turn types, with model-derived score differences correlating with sequence prevalence across conformations. However, these models do not fully capture the *β*-turn’s dynamic behavior, as residue pairs associated with static and dynamic states are not clearly separable in sequence likelihood space. This suggests that while sequence models encode relative conformational preference, the extent of conformational exchange depends on additional factors.

Our results also suggest that predicting, or designing, *β*-turn dynamics requires consideration of the adjacent structural context. Turns flanked by helices are strongly constrained and predominantly static, whereas strand- and coil-flanked turns display broader transition spectra. These observations indicate that *β*-turn behavior emerges from hierarchical constraints: local residue energetics define accessible basins, while the surrounding secondary-structure framework modulates transition pathways and kinetic accessibility. Collectively, our findings reposition *β*-turns from static chain reversals to dynamically interconverting micro-ensembles whose conformational landscapes are encoded by both sequence and context.

## Methods

### K-means Clustering

For each *β*-turn defined by consecutive residues *i* through *i* + 3, we construct the backbone dihedral vector

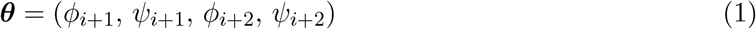

from the two central residues. This yields a dataset 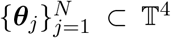 of *N* vectors on the four-dimensional torus with 2*π* periodicity. We seek *K* cluster centroids 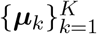that partition this dataset. For two dihedral vectors ***a*** = (*a*_1_, *a*_2_, *a*_3_, *a*_4_) ∈ 𝕋^4^, ***b*** = (*b*_1_, *b*_2_, *b*_3_, *b*_4_) ∈ 𝕋 ^4^, we define the toroidal distance

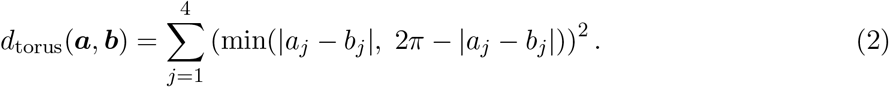

Each sample ***θ***_*j*_ is assigned to its nearest centroid via

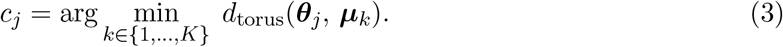

The centroids are updated with coordinate-wise circular means (i.e., averages computed with 2*π* wrap-around). Clustered points for different values of *K* are shown in Fig. S2.

To select the optimal number of clusters, we evaluated cluster quality using the Silhouette score over candidate counts *K* ∈ {4, 5, 6, 7, 8, 9}, running K-means ten times per *K* with different random initializations. For a sample ***θ***_*j*_ assigned to cluster *C*, the Silhouette score is

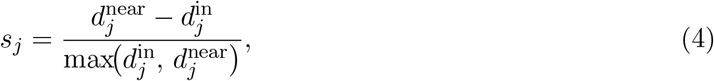

where 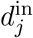 is the mean toroidal distance from sample *j* to all other members of its assigned cluster *C* (cohesion), and 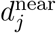 is the minimum, over clusters *C*′ ≠*C*, of the mean distance from sample *j* to members of *C*′ (separation from the nearest alternative cluster). A score of +1 indicates that a sample is well matched to its own cluster and poorly matched to its neighbors.

For each *K*, we report the average Silhouette score across all samples and initializations. As shown in Fig. S3, *K* = 6 attains the highest average Silhouette score and is chosen as the optimal cluster count. The training dataset consists of X-ray crystal structures sourced from the Protein Data Bank [8], specifically restricted to Escherichia coli proteins expressed in E. coli hosts.

### Identification and classification of *β*-turns from molecular dynamics trajectories

Molecular dynamics trajectories and corresponding protein topologies were obtained from the md-CATH database and analyzed using BioPython [20] (version 1.85) and MDTraj [21] (version 1.10.3). Structural features were extracted from the first frame of each trajectory, which served as the reference conformation. Backbone dihedral angles (*ϕ* and *ψ*) were computed for all residues, and secondary-structure assignments were determined using the DSSP algorithm [9].

*β*-turns were identified using a secondary-structure motif definition. Specifically, residues were selected if they matched the pattern [`-|H|E]TT[-|H|E`], where `T` denotes the two central residues of the turn and `H, E`, and `-` correspond to helix, *β*-strand, and coil assignments, respectively. This criterion captures *β*-turns occurring in diverse structural contexts while preserving a consistent local geometry.

Turn conformations were classified using the four backbone dihedral angles of the turn residues (*i* + 1, *i* + 2) via K-means clustering with *K* = 6, based on predefined centroids derived from the previous section. Representative examples were selected according to their distances to the cluster centroids in dihedral-angle space. Turns closest to each centroid were chosen as examples, and the corresponding backbone atom structures are shown in Fig. 1.

### Tracking *β*-turn dynamics across time

For each identified *β*-turn, turn types were assigned at every trajectory frame (1 ns resolution) by mapping the four backbone dihedral angles of residues *i* + 1 and *i* + 2 to the nearest centroid in dihedral-angle space. To ensure geometric consistency with a *β*-turn conformation, the C*α*–C*α* distance between residues *i* and *i* + 3 was monitored, and frames with distances exceeding 7.5 Å were excluded and labeled as non-turns. This procedure yielded a time-resolved classification of turn types for each trajectory.

To reduce noise arising from short-lived conformational fluctuations, turn-type assignments were temporally smoothed using a 10-frame persistence criterion. Only transitions lasting longer than 10 consecutive frames were retained; transient assignments were replaced by the modal turn type within a surrounding 19-frame window.

Using the smoothed annotations, we quantified turn-type transition counts and dwell times. Transition counts were calculated by summing all observed transition events between pairs of turn types. Dwell time was defined as the duration of a continuous interval spent in a given turn type before transitioning to a different type. These quantities were computed across all protein domains in the mdCATH dataset, with five independent replicas per domain weighted equally (1*/*5 each). Transition events and their associated durations were extracted from the trajectories and summarized as transition matrices.

### Analysis of alternative conformations in X-ray crystal structures

Protein structures containing alternate conformations were obtained from a curated catalog of crystallographic alternate locations (altlocs) [14]. *β*-turn motifs were identified from preassigned DSSP secondary-structure annotations in the catalog by selecting segments containing two consecutive turn-designated residues (`TT`). Segments containing three or more consecutive turn annotations were excluded to avoid ambiguous assignments. Turn types were then assigned by mapping the backbone dihedral angles (*ϕ* and *ψ*) of residues *i* + 1 and *i* + 2 to the previously derived K-means centroids.

For structures in which alternate conformations A and B corresponded to different *β*-turn types,the paired assignments were recorded in a correlation matrix (Fig. 2C–D) to quantify relationships between experimentally observed dual conformations. To reduce redundancy arising from closely related sequences, protein sequences were clustered using MMseqs2 [22], and each cluster was weighted equally in the statistical analysis. Representative examples of *β*-turns exhibiting dual conformations were visualized using UCSF ChimeraX [23] (version 1.9) and are shown in Fig. 2 E–F.

To complement crystallographic data, *β*-turn conformational variability was also analyzed using NMR ensembles. *β*-turn segments were identified from individual models within each NMR ensemble using the same DSSP-based motif [`H|E|-]TT[H|E|-`] and turn types assigned. For each *β*-turn, differences in assigned turn types across models within the same ensemble were recorded as correlated conformational pairs, analogous to altloc assignments in X-ray structures

### Amino acid preference analysis

Each *β*-turn was assigned to a single turn-type category based on its dominant conformation across time, defined as the turn type with the longest cumulative residence time over the trajectory. For each turn, amino acid identities at positions *i* + 1 and *i* + 2 were recorded, and pairwise amino acid counts were computed.

Within each turn-type category, turns were further stratified based on their dynamic behavior over the 500-ns trajectories. Turns that did not undergo any transitions (transition count = 0) were classified as static, whereas turns exhibiting at least three transition events were classified as dynamic. Turns with one or two transitions were excluded from this analysis to enhance separation between static and dynamic populations.

For each turn type, amino acid pair frequencies were calculated separately for static and dynamic turns. An enrichment score was defined as the difference between the frequency of a given amino acid pair in dynamic turns and its frequency in static turns. Positive scores indicate enrichment in dynamic turns, whereas negative scores indicate enrichment in static turns.

### Mutation and molecular dynamics simulation

Site-specific mutations were introduced by side-chain substitutions using the Dunbrack rotamer library [24] as implemented in ChimeraX (version 1.9) [25]. Rotamers were selected based on steric compatibility and local structural context prior to molecular dynamics (MD) simulations.

All MD simulations were performed using GROMACS (version 2025.2) [26–28]. The protein was modeled using the CHARMM22^∗^ force field [29]. The system was solvated in explicit TIP3P water in a cubic simulation box extending at least 1 nm from the protein surface, and Na^+^ and Cl^−^ ions were added to achieve electrostatic neutrality.

Energy minimization was first performed using steepest descent until convergence to remove steric clashes. Subsequently, equilibration was carried out in two stages under NPT conditions at 1 atm and 300 K. The first phase (10 ns) employed position restraints on heavy atoms to stabilize the solvent environment, followed by an unrestrained NPT phase (10−20 ns). Subsequently, the system was transitioned to an NVT equilibration step at 320K using Langevin stochastic dynamics implemented via the stochastic dynamics (sd) integrator, corresponding to a friction coefficient of ~ 0.1 ps^−1^ (*τ* ≈ 10 ps).

Production simulations were carried out in the NVT ensemble at 320 K using stochastic dynamics with a 2 fs integration timestep for a total of 250 million steps, corresponding to 500 ns of simulation time. Temperature was maintained at 320 K using Langevin dynamics (*τ* = 10 ps). Coordinates and energies were saved every 1 ns, with log data recorded every 100 ps. All covalent bonds involving hydrogen atoms were constrained using the SHAKE algorithm, enabling the larger timestep. Long-range electrostatics were treated using the particle mesh Ewald (PME) method with a 0.9 nm real-space cutoff. Van der Waals interactions employed the same cutoff with force-switch smoothing beginning at 0.75 nm. Neighbor searching was performed using the Verlet scheme with buffered cutoffs to ensure stable energy conservation.

### Conformational bias analysis using ProteinMPNN

Conformational bias between alternative *β*-turn states was evaluated using a structure-conditioned sequence scoring approach based on ProteinMPNN, following 18. Sequence preferences were compared across distinct backbone conformations via differences in model-assigned log-probabilities. Static *β*-turn segments were selected from the mdCATH dataset using the first frame of each trajectory as the reference conformation. For each example, an alternative backbone conformation was generated by guiding AlphaFold3’s reverse diffusion process from Gaussian noise to recover a structure **X** that maximizes a target likelihood ℒ [16, 30],

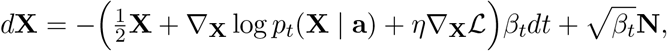

where **X** are atomic coordinates, **a** is the amino acid sequence, *β*_*t*_ is the noise schedule, and **N** ~ 𝒩 (**0, I**). The term ∇_**X**_ log *p*_*t*_(**X** | **a**) is the unconditional score function, while ∇_**X**_ℒ provides dihedral-angle guidance. The hyperparameter *η* = 0.1 scales the guidance term. We define the guidance objective

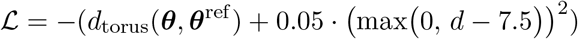

where 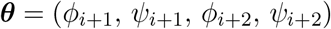 and 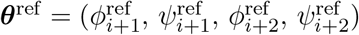are the predicted and target backbone dihedrals (in radians on 𝕋^4^, consistent with *d*_torus_ from Equation 2). Let 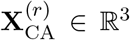 denote the C*α* coordinates of residue *r*, and 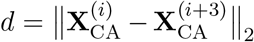. The second term is a one-sided quadratic penalty encouraging *d* ≤ 7.5 Å between the C*α* atoms of residues *i* and *i* + 3. Generated structures are relaxed with the AMBER99 force field [31] to alleviate steric clashes.

This procedure generated pairs of backbone conformations corresponding to different *β*-turn types for the same sequence context. ProteinMPNN [19] was used to evaluate sequence likelihoods conditioned on backbone geometry. For each conformation, the log-probability of the amino acids at positions *i* +1 and *i* +2 was computed with the backbone fixed. Conformational bias was defined as.

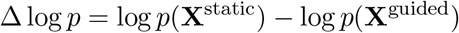

using ProteinMPNN scores for the residue pair under guided **X**^guided^ and static **X**^static^ conformations. To relate sequence-based bias to dynamics, Δ log *p* was compared with the log_2_ prevalence ratio of residue pairs from molecular dynamics trajectories (Fig. 4), with pairs grouped by dynamic/static enrichment (Fig. 5). Pearson correlations were computed over residue pairs with at least 100 observations in either conformation to reduce noise. Finally, molecular dynamics simulations were initialized from alternative backbone conformations for selected sequences. The resulting trajectories were analyzed to assess whether predicted conformational bias corresponded to differences in dynamic behavior.

## Data Availability

Code available at: https://doi.org/10.5281/zenodo.19891630, data available at: https://doi.org/10.5281/zenodo.19890778

## Supplementary information

**Figure S1:**
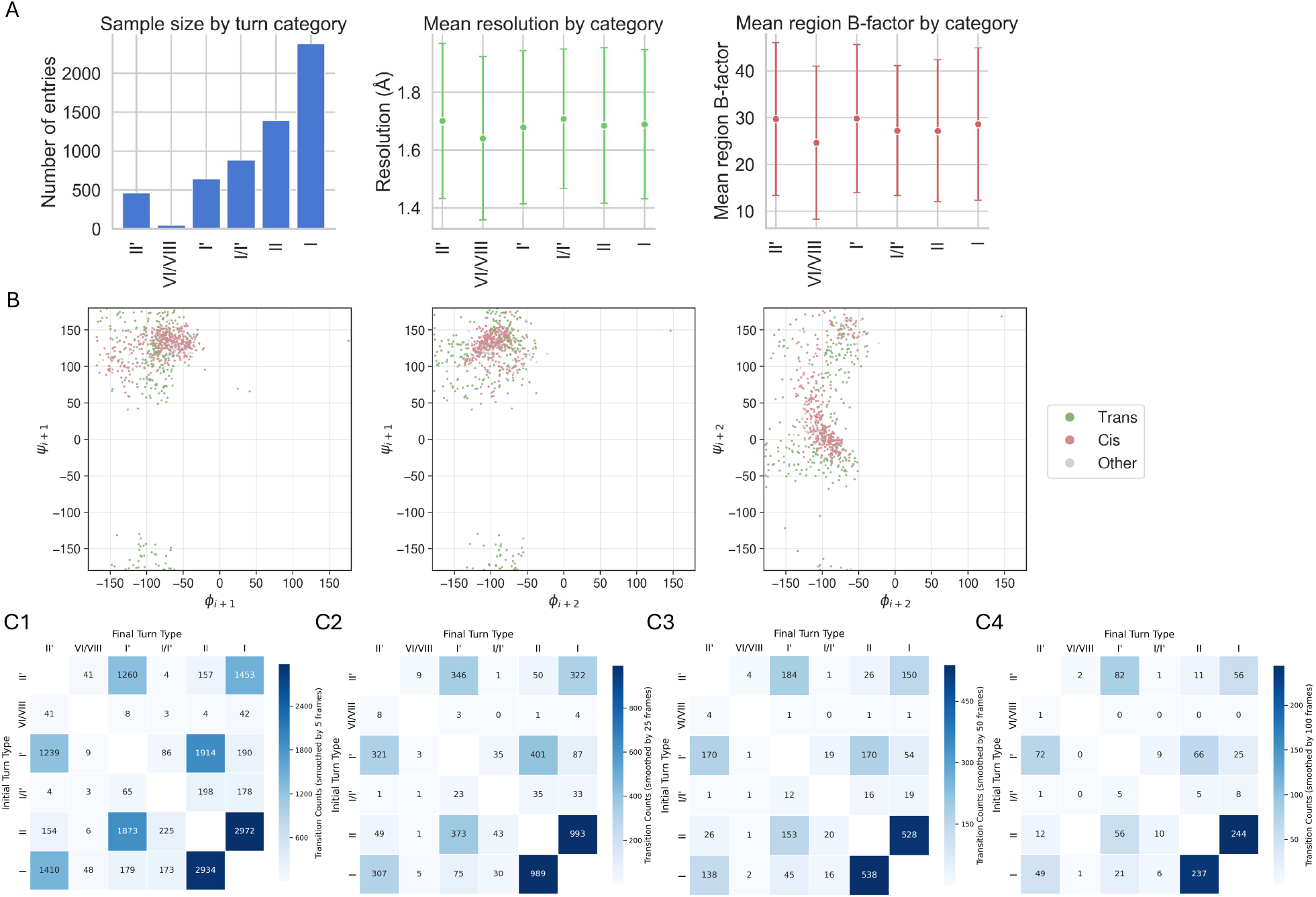
Validation of clustering quality, structural interpretation, and robustness of *β*-turn dynamic analysis. **(A)** Sample size (left), mean crystallographic resolution (middle), and mean regional B factor (right) for each *β*-turn type. Similar distributions across turn types indicate that clustering is not biased by structural quality or local flexibility of the underlying experimental data. **(B)** Cis and trans conformations within the `VI/VIII` cluster visualized in Ramachandran representations. Points are colored by peptide-bond configuration, showing that cis (pink) and trans (green) turns occupy overlapping regions in all three dihedral projections, supporting the combined classification of `VI` (cis) and `VIII` (trans) into a single cluster. **(C)** Transition count matrices computed using different temporal smoothing windows: 5 frames *(C1)*, 10 frames (Fig. 2A), 25 frames *(C2)*, 50 frames *(C3)*, and 100 frames *(C4)*. Overall transition patterns between *β*-turn types are preserved across smoothing parameters, demonstrating robustness to the choice of smoothing.

**Figure S2:**
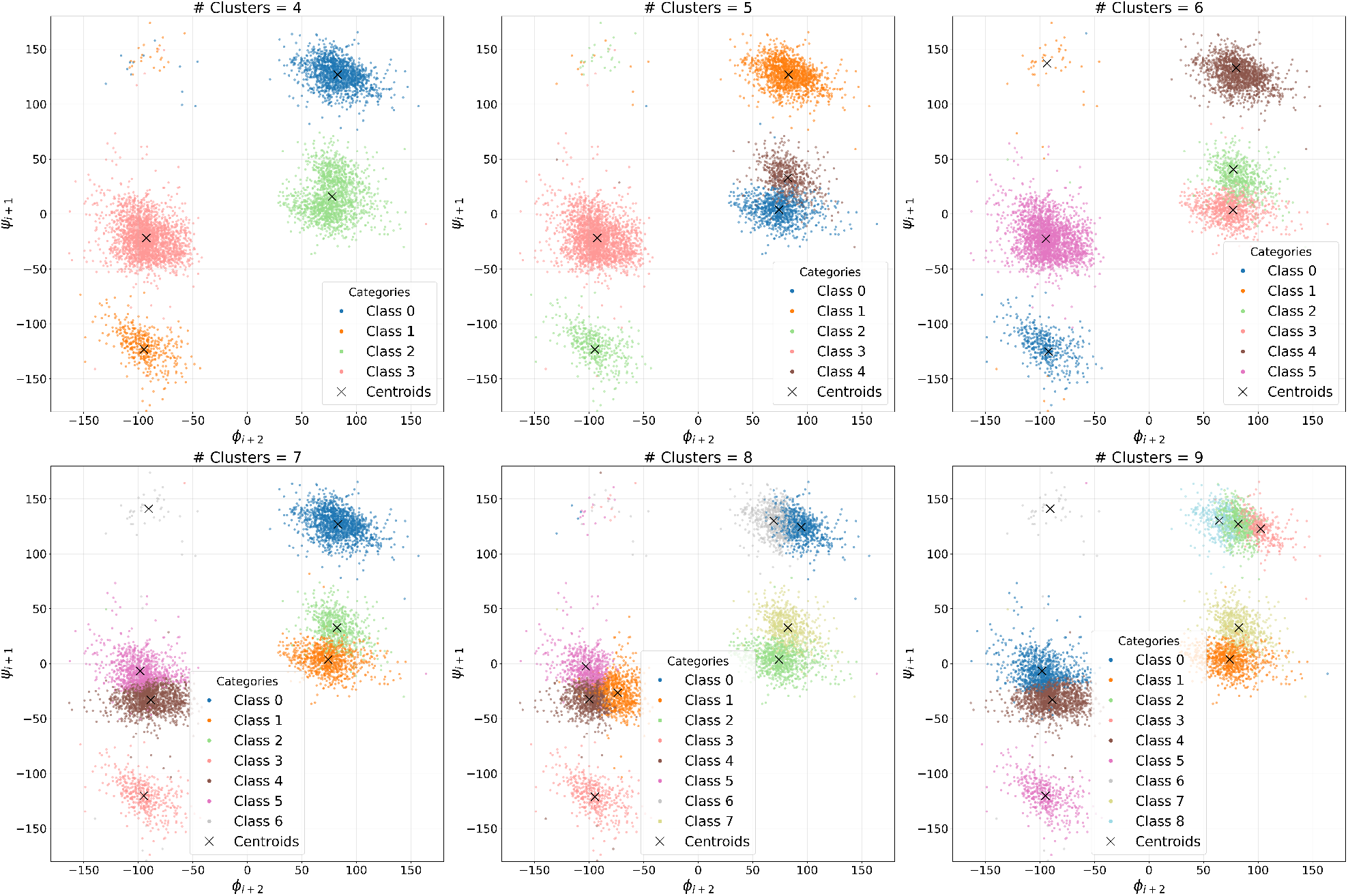
Cross-bond plot of dihedral angles *ϕ*_*i*+2_ and *ψ*_*i*+1_ clustered by K-means with different numbers of clusters.

**Figure S3:**
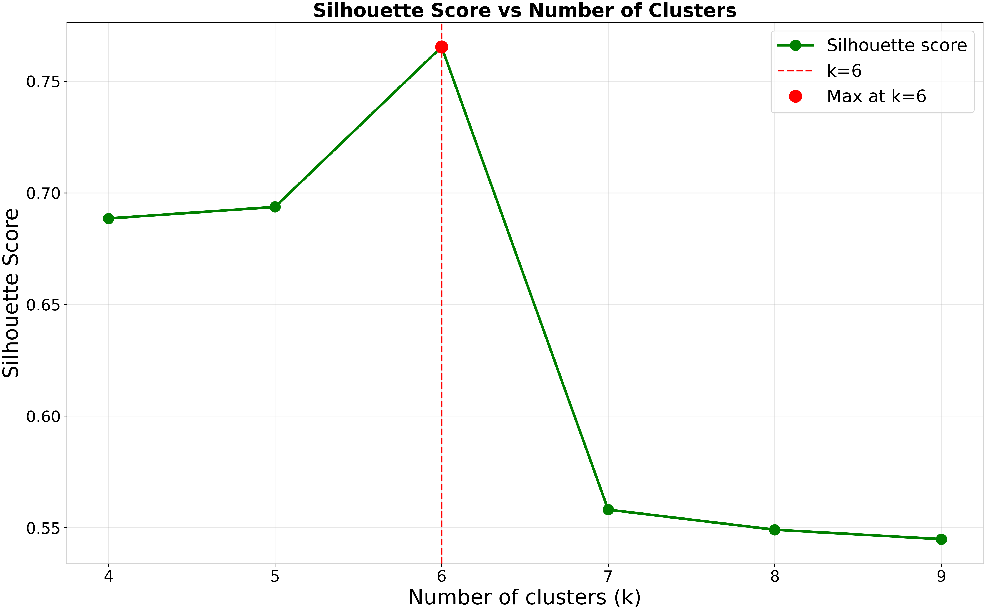
Silhouette scores are compared across K-means runs with varying numbers of clusters.

**Figure S4:**
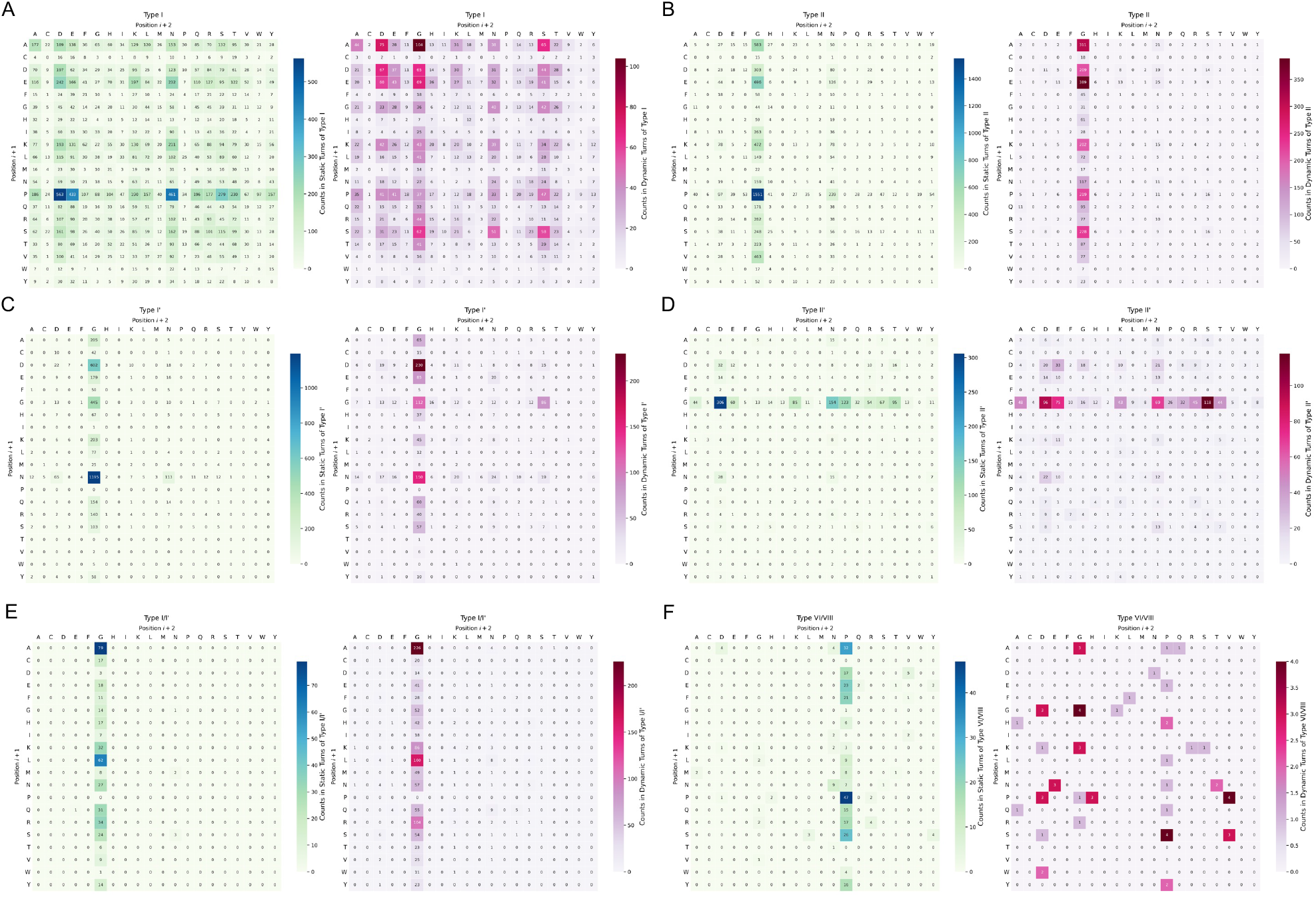
Pairwise amino acid composition of static and dynamic subclasses within each *β*-turn type. For each turn type, residue-pair frequencies at positions *i* + 1 and *i* + 2 are shown separately for static (left; green-to-blue gradient) and dynamic (right; purple-to-red gradient) subclasses. Values represent raw occurrence counts and correspond to the enrichment analysis presented in the main text. **(A)** In type `I` turns, residue pairs containing `PRO` at position *i* + 1 (e.g., `PRO-ASP, PRO-ASN, PRO-GLU`) are concentrated in static turns, whereas pairs with `GLY` at position *i* + 2 or `ALA` at position *i* + 1 (e.g., ALA-GLY, GLY-GLY, ALA-ASP, SER-GLY) are more prevalent in dynamic turns. **(B)** Type `II` turns show strong accumulation of the `PRO-GLY` pair in static turns, while `ALA-GLY, GLU-GLY, SER-GLY`, and `ASP-GLY` are more frequent in dynamic turns. **(C)** In type `I′` turns, `ASN-GLY` and `GLY-GLY` are concentrated in the static subclass, whereas `GLY-SER` occurs primarily in dynamic turns; `ASP-GLY` shows comparable representation in both subclasses. **(D)** Type `II′` turns show higher counts of `GLY-ASP` and `GLY-ASN` pairs in the static subclass, whereas `GLY-SER` is enriched in dynamic turns. **(E)** The hybrid `I/I′` type exhibits an overall dynamic tendency, with higher counts of several residue pairs in the dynamic subclass, particularly `ALA-GLY` and `LEU-GLY`. **(F)** Type `VI/VIII` turns are predominantly static, with strong accumulation of `ALA-PRO` and `PRO-PRO` pairs in the static subclass; dynamic examples are sparse, consistent with the overall static tendency of this type.

**Table S1:** Example for each *β*-turn type shown in Figure 1B.

| PDB ID | Residue | Sequence | $\phi_{i+1}$ | $\psi_{i+1}$ | $\phi_{i+2}$ | $\psi_{i+2}$ | Type |
| --- | --- | --- | --- | --- | --- | --- | --- |
| 6IAH | 178–181 | LKPD | −66.89 | 138.05 | −104.62 | 34.16 | VI/VIII |
| 3SFV | 342–345 | NKDQ | −65.04 | −19.00 | −102.19 | 3.96 | I |
| 3JU4 | 636–639 | VGDD | 51.59 | −118.78 | −94.60 | 2.91 | II′ |
| 2XVS | 332–335 | NSGA | −55.74 | 129.90 | 80.34 | 1.11 | II |
| 3HDJ | 70–73 | IKGQ | 45.79 | 41.59 | 79.85 | 0.40 | I′ |
| 6A02 | 131–134 | LAGV | −87.73 | −0.21 | 81.73 | 22.60 | I/I′ |

**Table S2:**
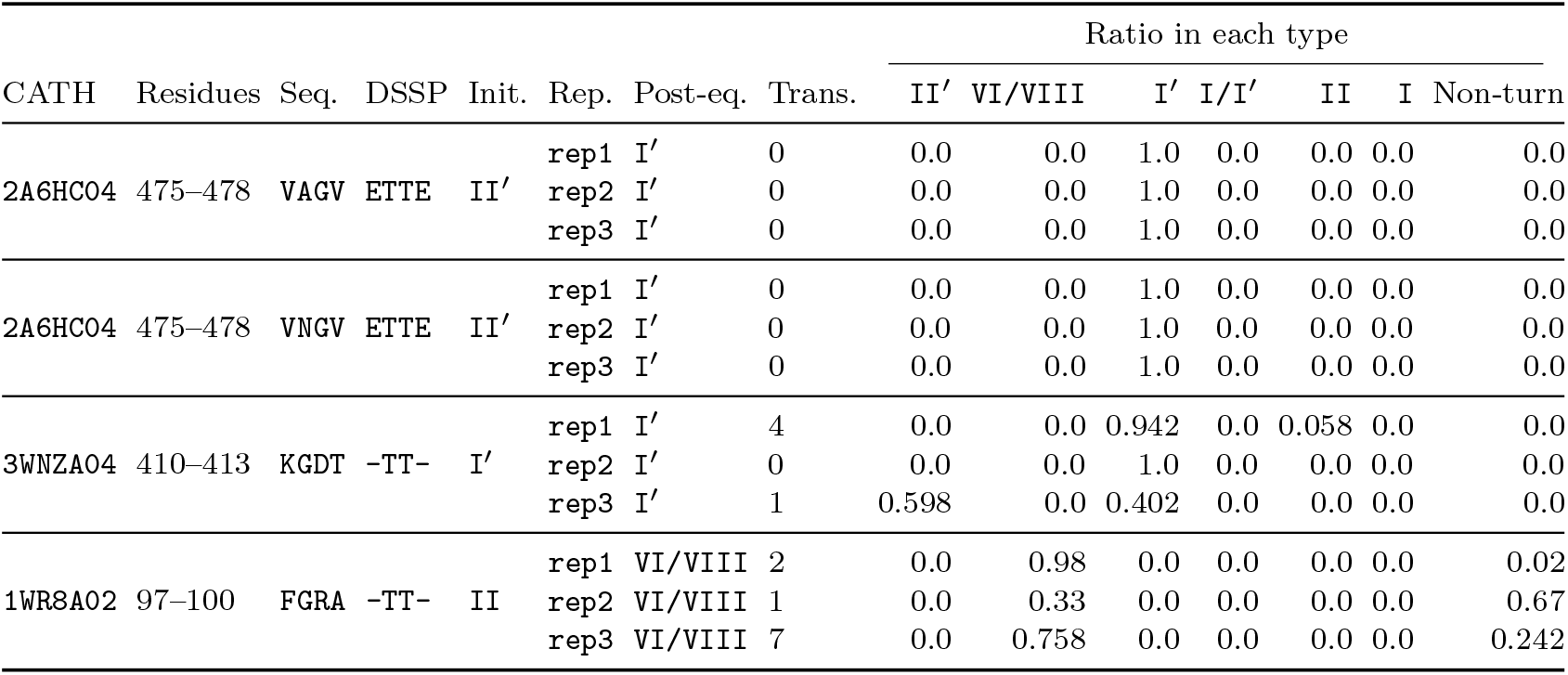
ProteinMPNN-predicted conformational bias and observed *β*-turn dynamics across replicate simulations. The table summarizes *β*-turn behavior for selected protein systems initialized in specific conformations and simulated in three independent replicas.

**Table S3:**
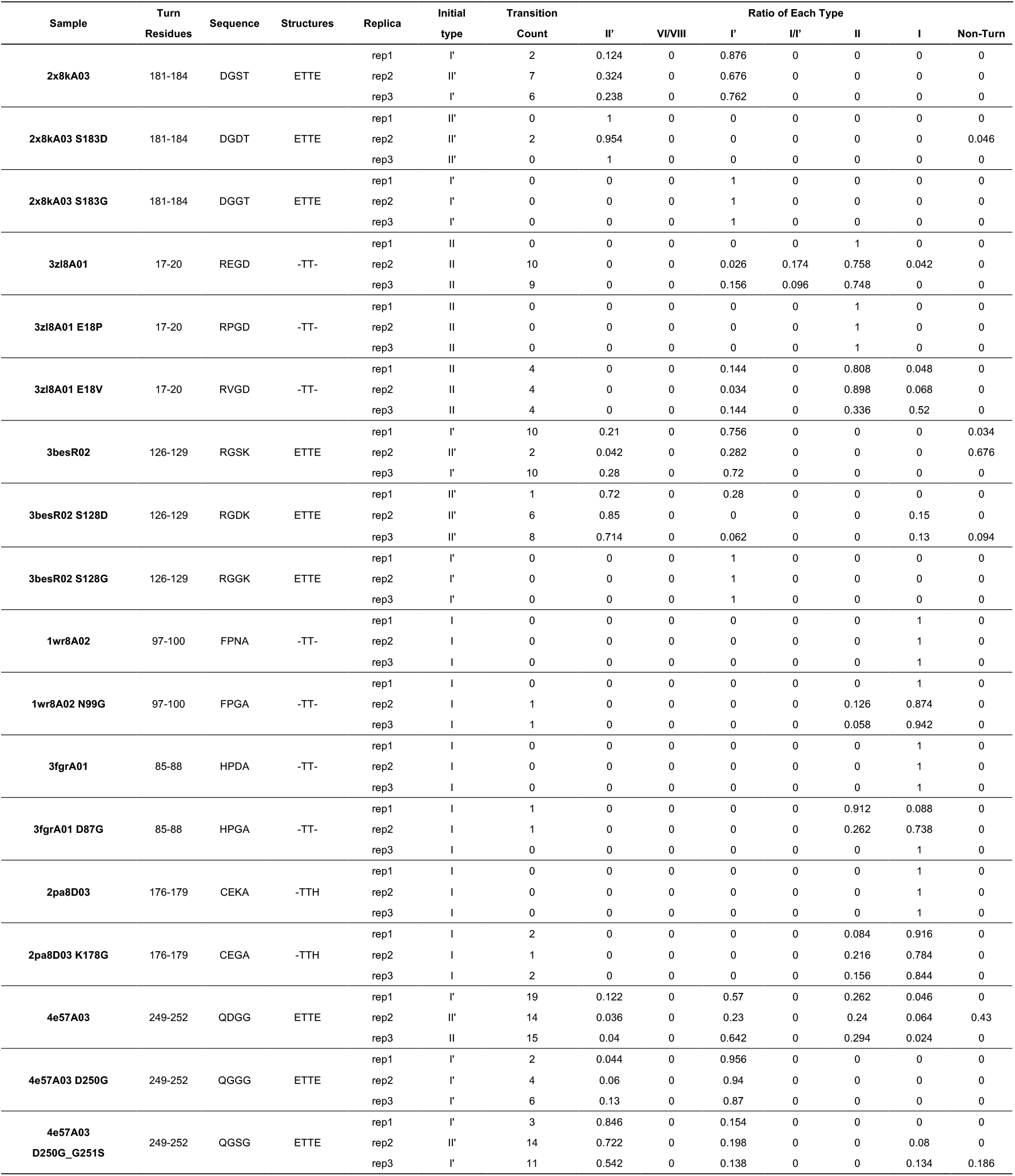
Summary of *β*-turn dynamics across replicate trajectories. The table summarizes turn behavior for each protein system (wild type and mutants) simulated in three independent replicas under identical conditions. “Sample” denotes the CATH identifier and mutation. “Turn residues” indicate the *β*-turn segment. “Sequence” and “Structures” list the amino acid sequence and DSSP annotation of the turn (positions *i* to *i* + 3). “Initial turn type” corresponds to the conformation assigned at the first trajectory frame. “Ratio in each type” represents the fraction of frames classified as a given turn type relative to the total length of trajectories.

